# Barley is a potential trap crop for root parasitic weeds broomrapes

**DOI:** 10.1101/2024.10.22.619574

**Authors:** Maiko Inoue, Xiaonan Xie, Kaori Yoneyama

## Abstract

Root parasitic weeds broomrapes (*Phelipanche* and *Orobanche* spp.) cause devastating damages to agricultural production all over the world. The seeds of broomrapes germinate when they are exposed to germination stimulants, mainly strigolactones (SLs), released from roots of any plant species, while broomrapes parasitize only dicot plants. Therefore, monocots can be trap crops for broomrapes, as they induce seed germination but are not parasitized. In this study, we examined two European and one Japanese cultivars of barley for their potentials as trap crops for broomrapes and found that the European cultivars, Sebastian and Golden promise, are good potential trap crops as they produce more SLs and exhibit higher mycorrhizal colonization rates as compared to the Japanese cultivar Shunrai.

## Introduction

The root parasitic weeds broomrapes (*Phelipanche* and *Orobanche* spp.) are causing devastating damages to agricultural production all over the world. They are holoparasites lacking photosynthetic activity and depend entirely on their hosts for the supply of water and nutrients. Areas threatened by broomrapes are 16 million ha in the Mediterranean and west Asia with a negative impact on yields of 20%-100%.^1^)

The tiny seeds of broomrapes have limited energy stock which allows germination and elongation of radicles only to a few millimeters, and the germinated seeds need to parasitize host roots within a few days or they will die. To avoid germination in the absence of hosts in the vicinity, parasite seeds germinate only when they are exposed to germination stimulants released from plant roots. Strigolactones (SLs) are the most important and widely distributed germination stimulants, and so far, more than 40 SLs have been characterized from root exudates of various plant species.^2)^

Plants exude SLs not for root parasites but for arbuscular mycorrhizal (AM) fungi to commence symbiosis.^3)^ More than 80% of land plants form symbiotic relationships with AM fungi in which AM fungi supply mineral nutrients such as nitrogen and phosphorous to host plants and in turn receive photosynthates from the host. In plants, SLs function as a hormone inhibiting shoot branching.^4,5)^ Biosynthetic mutants defective in SLs exhibit highly branched phenotype and exogenous SL analogues restore the phenotype. SL deficit mutants are resistant to root parasitic weeds as they induce less germination, but, at the same time, they have lower levels of AM colonization.^4,6)^ Therefore, in general, these mutants are less efficient in nutrient acquisition and more susceptible to biotic and abiotic stresses.^7^)

One of approved strategies to mitigate the damages caused by root parasitic weeds is to grow ‘trap crops’; non-host crops induce seed germination of root parasitic weeds.^8-10)^ In the case of witchweeds (*Striga hermonthica* and *S. asiatica*) parasitizing monocots, cotton and legumes including soybean, groundnut, and cowpea, have been successfully used as trap crops.^11)^ Since these trap crops induce *Striga* germination but not serve as their hosts, trap crops eventually reduce seed bank of *Striga*. For broomrapes, monocots can be trap crops and so far wheat and maize have been examined for their effects on seed germination stimulation and parasitism of broomrapes.^10,12)^ SLs produced by wheat and maize have been well characterized.^13-15)^ Especially in maize cultivars, changes in SL profile have been shown to contribute to resistance to *Striga*.^16)^

Barley (*Hordeum vulgare*) is the fourth most important cereal crop of the world after wheat, corn, and rice.^7)^ Barley was first domesticated in southwestern Asia and is now widely distributed in relatively cool areas such as Russia, Canada, and many European countries where broomrapes are causing severe damages to dicot crops. Major SLs of wheat and maize are orobanchol and zealactone-related compounds, respectively, SLs of barley have been characterized only recently as 6-*epi*-heliolactone^17)^ and two novel barley SLs (tentatively named barleylactones, BL1 and BL2).^18)^ Structural determination of these novel SLs will be reported elsewhere. Germination stimulation activities of BL1 and BL2 to broomrape seeds are shown in Figure S1.

In the present study, we examined two European (Sebastian and Golden Promise) cultivars and one Japanese cultivar (Shunrai) of barley for their potentials as trap crops for broomrapes by comparing their SL productions, germination stimulation activities of root exudates on branched broomrape (*Phelipanche ramosa*) and clover broomrape (*Orobanche minor*), and rates of AM colonization. In addition, effects of nutrient deficiency on SL exudation and shoot branching in the barley cultivar Shunrai were evaluated.

## Materials and methods

### 1. Seeds

Barley seeds of cultivar Shunrai were obtained from a local shop in Japan. Cultivars of Golden Promise and Sebastian were kindly provided by Dr. Phillip Brewer (Adelaide University, Australia) and Dr. Marek Marzec (University of Silesia, Poland), respectively. *Orobanche minor* seeds were collected from mature plants that parasitized red clover (*Trifolium pratense* L.) grown in Tochigi Prefecture, Japan in June 2020. *Phelipanche ramosa* (pathotype 1) seeds collected from mature plants parasitizing oilseed rape plants were supplied by Dr. Jean-Bernard Pouvreau (University of Nates, France).

### 2. Barley growth conditions

Barley plants were grown in pots (volume, 700 mL; diameter, 12.4 cm; depth, 13.3 cm) filled with vermiculites and compost (1 : 1 v/v) under 16 h light (approximately 240 µmol m^−2^ s^−1^) / 8 h dark at 23°C. They were watered when necessary.

### 3. Collecting root exudates

Tap water (100–200 mL) was poured onto the soil surface and root exudates eluted from the holes of pot bottom were collected, which were extracted with ethyl acetate. The ethyl acetate phase was dried over anhydrous MgSO_4_ and concentrated *in vacuo*. All crude samples were stored at 4°C until use.

### 4. SL analysis

We analyzed barley SLs by using ultra performance liquid chromatography coupled to tandem mass spectrometry (UPLC-MS/MS). For this, the Acquity UPLC System (Waters) coupled to a Xevo TQD triple-quadrupole mass spectrometer (Waters MS Technologies) with electrospray (ESI) interface was used.

Chromatographic separation was achieved using an ODS column (ACQUITY UPLC, BEH C_18_, 2.1 × 100 mm, 1.7 µm; Waters) with a water-MeOH gradient containing 4% 50 mM ammonium acetate to promote ionization. Separation started at 35% MeOH, followed by a 2 min gradient to 55% MeOH, followed by a 13 min gradient to 95%, kept 96% MeOH for 2 min to wash column and then back to 35% MeOH for 3 min. The column was equilibrated at this solvent composition for 5 min before the next run. Total runs time was 25 min. The column oven temperature was maintained at 40°C with a flow-rate of 0.2 mL min^−1^ (sample injection volume of 1 µL). MRM transitions for 6-*epi*-heliolactone eluting at 5.5 min were monitored for *m/z* 361/97 at a collision energy (CE) of 18 V and *m/z* 361/233 at CE of 18 V with a cone voltage of 25 V. The MRM transitions of *m/z* 377/97 at CE of 20 V and *m/z* 377/231 at CE of 15 V with a cone voltage of 25 V were used for the detection of BL2 eluting at 4.4 min.

### 5. Germination assay

Germination assays with *O. minor* and *P. ramosa* seeds were conducted in a manner similar to that reported previously.^19)^ In short, an aliquot of either root exudate samples corresponding to 32 µL to 20 mL of root exudates (100 mL) in ethyl acetate or acetone solution of the synthetic SL, GR24 (mixture of four stereoisomers, 10^−6^ M), as positive control was added to a 5-cm Petri dish lined with a filter paper. The solvent was allowed to evaporate before the discs carrying the seeds conditioned at 23°C for 7 days were placed on the filter paper and treated with sterile Milli-Q water (650 µL). The Petri dishes were sealed, enclosed in polyethylene bags, and placed in the dark at 23°C for 5 days. Seeds were considered germinated when the radicle protruded through the seed coat.

### 6. Mycorrhizal colonization

To determine the mycorrhizal colonization, each barley plant was grown in pots (volume, 700 mL; diameter, 12.4 cm; depth, 13.3 cm) filled with vermiculites containing mycorrhizal materials (*Rhizophagus* sp.). Plants were subjected to a complete phosphate deficiency and roots were harvested on 28 days after sowing. Collected roots were soaked in 10% KOH aqueous solution and heated at 80°C for 30 min, and KOH was removed, and replaced with 0.05% w/v trypan blue in lactoglycerol. Roots in trypan blue solution were heated 80°C for 30 min again. After staining, roots were washed with LG solution 80°C for 30 min. The vesicle formation rate was determined by the lattice intersection as the percentage of cystic states present at the intersection.^20^)

### 7. Effects of nutrient deficiency on SL exudation and shoot branching

As the seeds of the European cultivars were limited, we used Japanese cultivar Shunrai in this experiment. Barley plants (cv. Shunrai) were grown in hydroponics and effects of phosphate and nitrogen deficiency on SL exudation and shoot branching were examined. Plants were grown in tap water for 7 days after sowing and grown in 1/2TT media^21)^ for another 7 days. Then, plants were subjected to each nutrient condition for 14 days. Collected root exudates were extracted with ethyl acetate and SLs were analyzed by LC-MS/MS.

## Results

### 1. SL levels in root exudates collected during 28–44 days after sowing

Root exudates collected on 28, 34, 39, and 44 days after sowing were analyzed by LC-MS/MS for their SLs, 6-*epi*-heliolactone, BL1 and BL2 (Fig. 1). In all root exudate samples collected, levels of BL1 were below the detection limit of our LC-MS/MS. Levels of both 6-*epi*-heliolactone and BL2 increased with cultivation period mainly due to the increase in root biomass. Among the three cultivars, Golden Promise appeared to release highest levels of SLs, and the Japanese cultivar Shunrai was a poor SL releaser as compared to the European cultivars (Fig. 2).

**Fig. 1.**
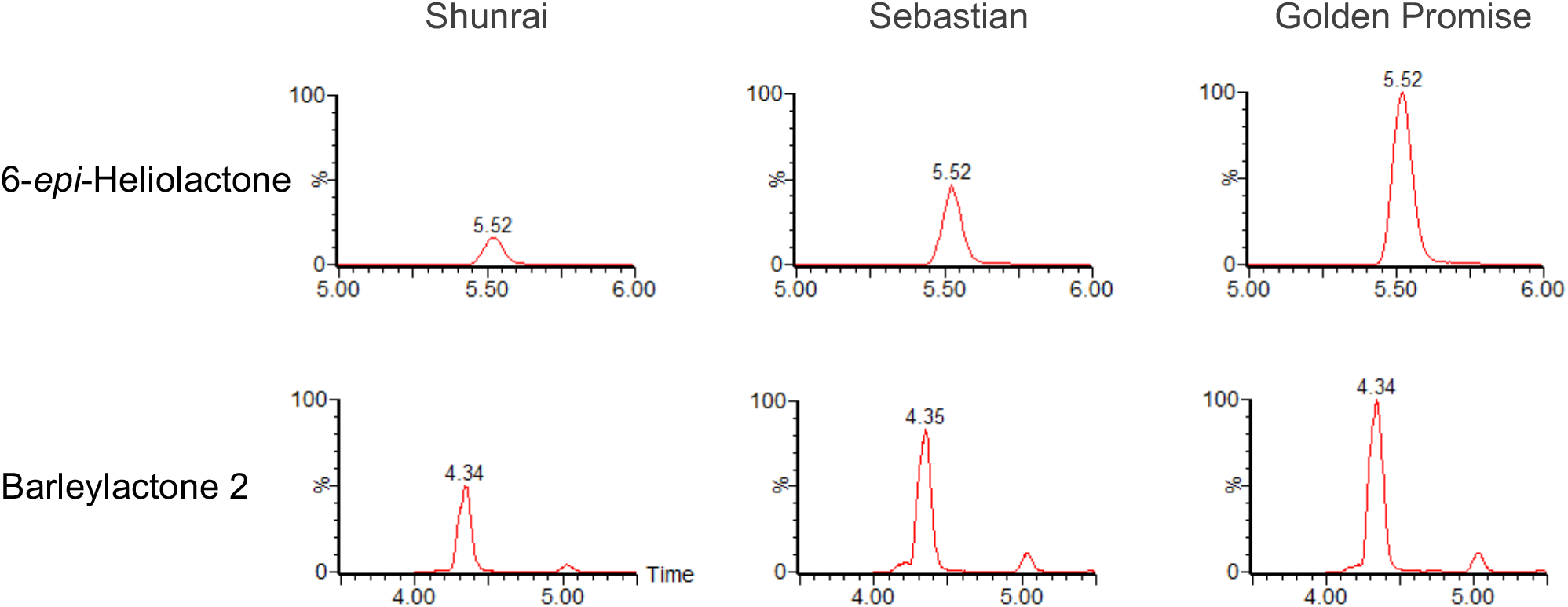
Identification of SLs. MRM chromatograms of 6-*epi*-heliolactone (*m/z* 361.0/97.0) and barleylactone 2 (*m/z* 377.0/97.0) in root exudate extracts of Shunrai, Sebastian and Golden Promise are shown (ESI-MS positive mode).

**Fig. 2.**
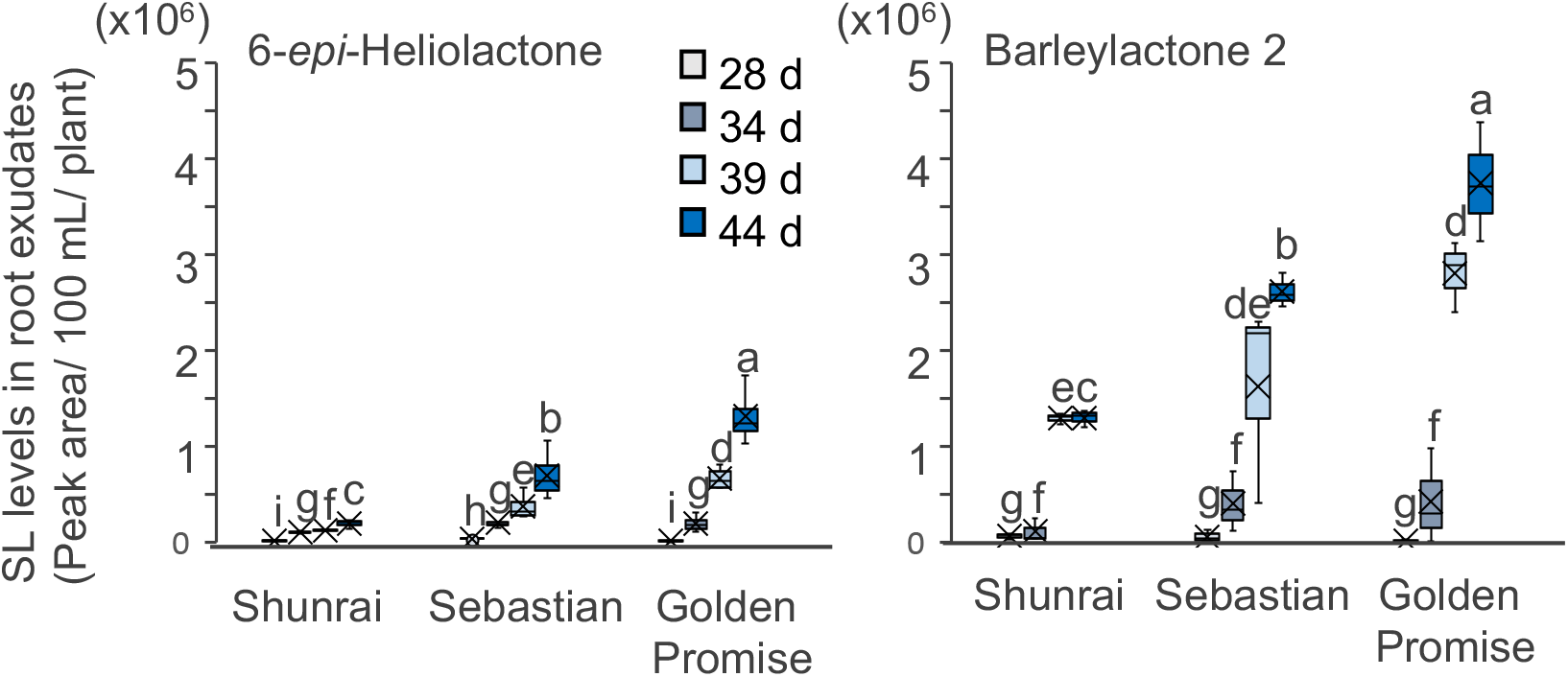
SL levels in barley root exudates. Plants were grown in vermiculites and root exudates were collected on 28 (light gray box), 34 (gray box), 39 (light blue box), 44 (blue) days after sowing. n=3. The different letters denote a statistically significant difference at *P*<0.05 among different cultivars according to the ANOVA, Tukey-Kramer HSD test.

### 2. Germination stimulation activity of root exudates

Germination stimulation activities of root exudates collected on 44 days after sowing from three barley cultivars on *O. minor* and *P. ramosa* seeds were examined. No or negligible germination was observed for the seeds incubated in Milli-Q water, negative control (data not shown). The synthetic SL, GR24, induced >60% germination of *O. minor* and *P. ramosa* seeds and thus both seeds are equally sensitive to GR24.

In general, root exudates from the European cultivars Sebastian and Golden Promise were more active than that of Shunrai in germination stimulation of both *O. minor* and *P. ramosa* seeds. The seeds of *O. minor* appeared to be slightly less sensitive to root exudates of barley than those of *P. ramosa*. Germination stimulation activities of root exudates on *P. ramosa* increased in a dose dependent manner while they were active only at high doses in *O. minor* germination (Fig. 3).

**Fig. 3.**
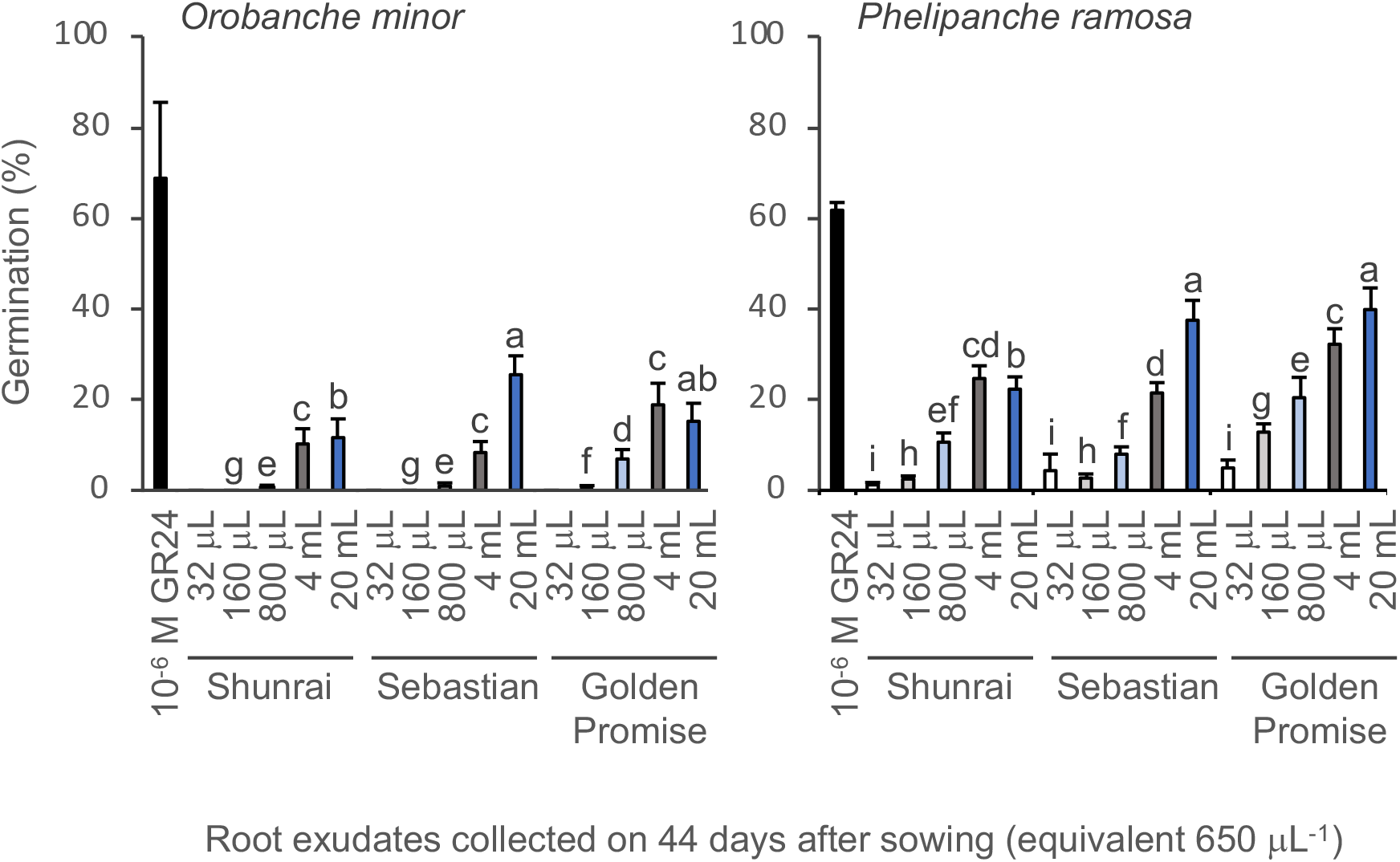
Germination stimulation activities of barley root exudates to broomrapes. The root exudate samples collected on 44 days after sowing were used in germination assay. Aliquots of ethyl acetate solutions of root exudate samples corresponding to 32, 160, 800 mL, 4 and 20 mL of media were to transferred to 5 cm Petri dishes lined with a filter paper. *n* = 9-12. The different letters denote a statistically significant difference at *P*<0.05 among different cultivars according to the ANOVA, Tukey-Kramer HSD test.

### 3. Mycorrhizal colonization

Mycorrhizal colonization rates were compared among Shunrai, Sebastian, and Golden Promise grown in pots under nutrient starved conditions. The highest and lowest mycorrhizal colonization rate was found in Sebastian and Shunrai, respectively under our experimental conditions (Fig. 4).

**Fig. 4.**
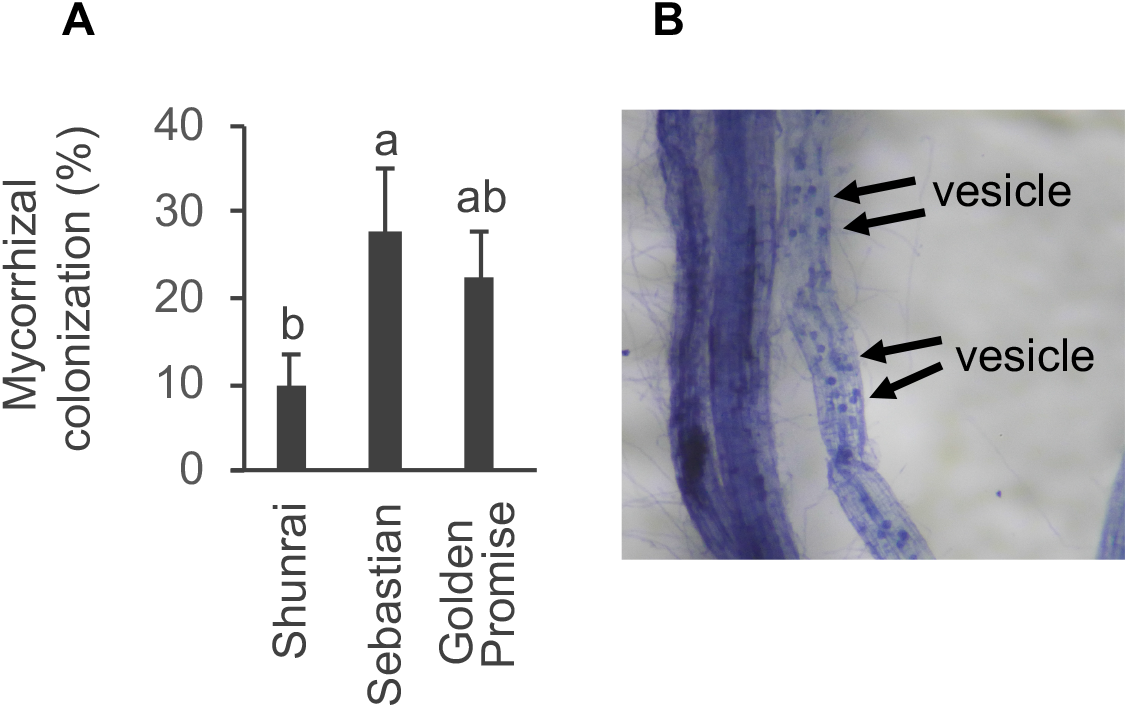
Mycorrhizal colonization rates in barley roots harvested on 28 days after sowing. (A) Mycorrhizal colonization rates. (B) Mycorrhizal roots dyed with TB. The different letters denote a statistically significant difference at *P* < 0.05 among different cultivars according to the ANOVA, Tukey-Kramer HSD test. *n* = 5.

### 4. Shoot branching phenotypes

Three barley cultivars, Shunrai, Sebastian, and Golden Promise were grown in pots and the number of shoot branching (tillers) was determined on 28 and 44 days after sowing. The number of shoot branching was counted as one which the tiller was over 1 cm (Fig. 5). Although no differences in the number of shoot branching were observed on 28 days, Sebastian and Golden Promise showed statistically significant more tillers than Shunrai on 44 days after sowing under our experimental conditions. Shunrai did not increase tillers during the period from 28 to 44 days after sowing, while Sebastian and Golden Promise achieved two-fold increase in the number of tillers in this period.

**Fig. 5.**
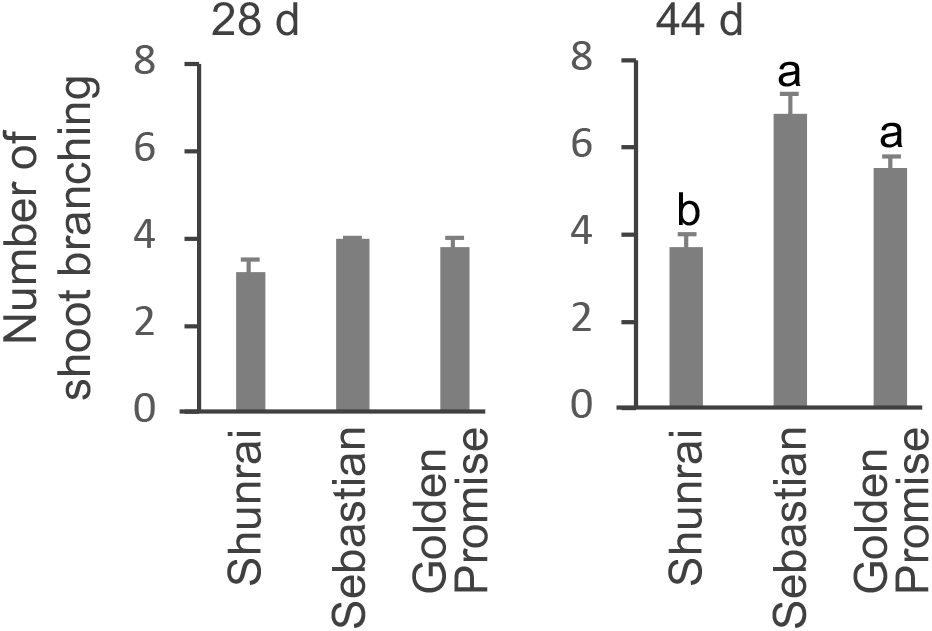
Shoot branching phenotypes of three barley cultivars on 28 and 44 days after sowing. Number of shoot branching (*n* = 3–12). The different letters denote a statistically significant difference at *P* < 0.05 among different cultivars according to the ANOVA, Tukey-Kramer HSD test.

### 5. Effects of nutrient deficiency on shoot branching, plant biomass, and SL exudation

Effects of nitrogen and phosphate deficiency on shoot branching, plant biomass, and exudation of 6-*epi*-heliolactone and BL2 were examined with Japanese cultivar Shunrai. Phosphate and nitrogen deficiency severely suppressed shoot branching (Fig. 6A) and resulted in the reduced shoot biomass but did not greatly affect root biomass (Fig. 6B). SLs in root exudates were below the detection limit when barley plants were grown under normal nutrient conditions, and nutrient deficiency, especially nitrogen deficiency, significantly promoted SL exudation (Fig. 6C).

**Fig. 6.**
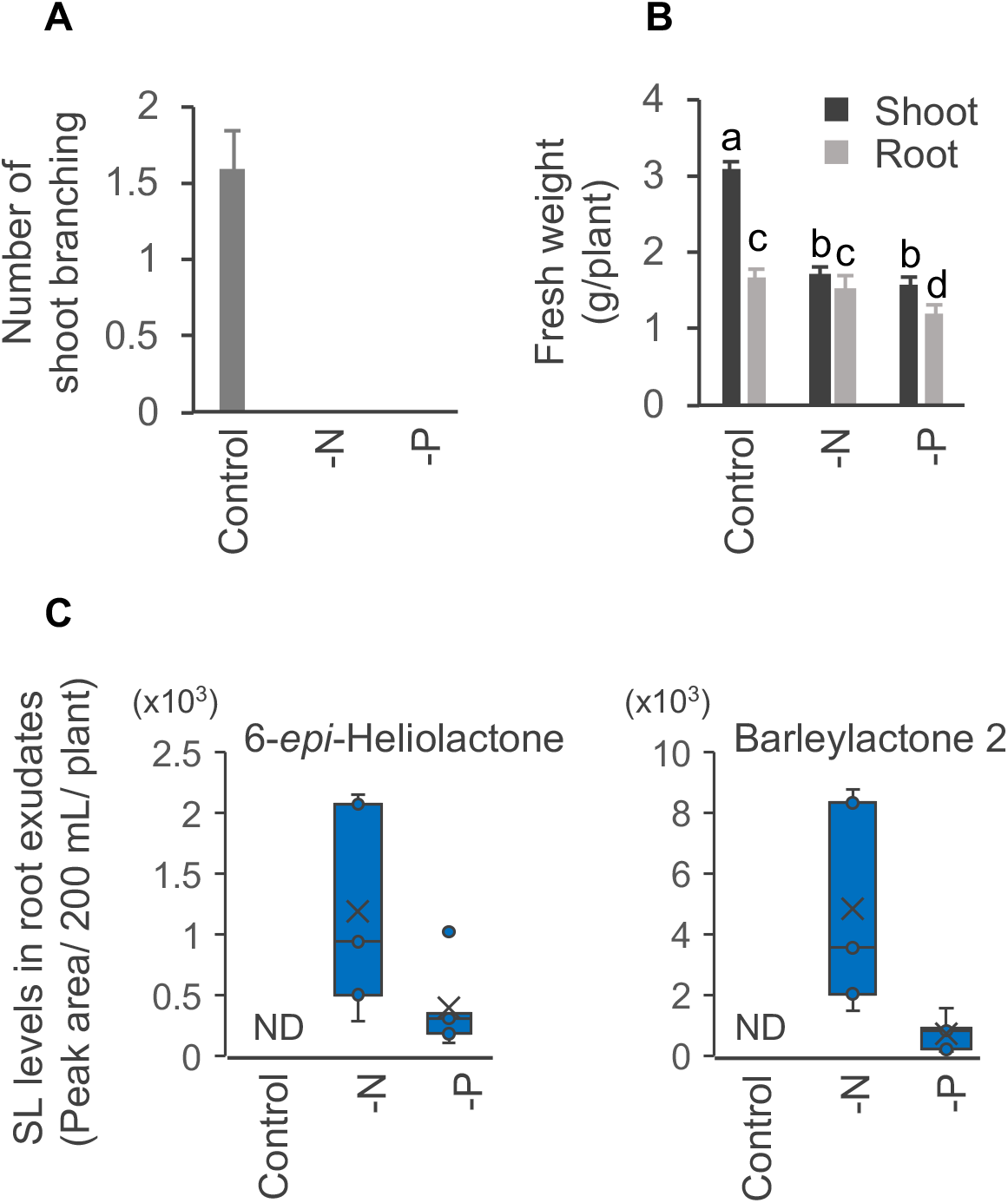
Effects of nutrient deficiency in Shunrai. (A) Shoot branching (B) Shoot and root fresh weight (C) and SL exudation. The different letters denote a statistically significant difference at *P* < 0.05 according to the ANOVA, Tukey-Kramer HSD test. *n* = 3-5.

## Discussion

Barley plants exude at least four SLs including 6-*epi*-heliolactone, which had been isolated from black oat (*Avena strigosa*),^17)^ and novel SL, BL2.^18)^ However, under our experimental conditions, levels of other barley SLs were below the detection limit of LC-MS/MS analysis and therefore 6-*epi*-heliolactone and BL2 were estimated based on their peak areas. Known canonical SLs could not be detected from root exudates of the three barley cultivars in our experimental conditions, though it was already reported that barley exudes 5-deoxystrigol.^22)^ We cannot exclude the possibility that barley exudes minute amounts of 5-deoxystrigol. Different growth conditions may also affect the profile of SLs. Although MRM analysis is a good method for quantification of SLs, it is important to confirm SL characterization by comparing product ion scan data with those of SL standards.

The three barley cultivars produced similar levels of SLs in the early growth stage (28–34 days after sowing), and then differences in SL levels became evident in the later growth stage (39–44 days after sowing); SL levels in root exudates increased in the order Shunrai < Sebastian < Golden Promise. According to the SL levels estimated based on their peak areas in LC-MS/MS, levels of BL2 appeared to be higher than those of 6-*epi*-heliolactone in root exudates of these barely cultivars, suggesting that BL2 may be more important than 6-*epi*-heliolactone in AM symbiosis of barley. SL levels in root exudates of the barley cultivars seem to be positively correlated with AM colonization rates as Shunrai, the poor SL producer, exhibited the lowest colonization rate (Fig. 4). However, as SLs function only in the pre-symbiotic stage, other factors also affect AM symbiosis. For example, Sebastian exudes less SLs than does Golden Promise, AM colonization rate was slightly higher in Sebastian than in Golden Promise.

In the case of grapevine rootstocks, 6-*epi*-heliolactone was suggested to be more important than the other non-canonical SL, vitislactone, in AM colonization.^23)^ As BL2 seems to be less stable than 6-*epi*-heliolactone,^18)^ 6-*epi*-heliolactone would contribute more to the promotion of AM colonization in barley as well. However, BL2 would stay longer in root exudates and also in the slightly acidic rhizosphere enough to exhibit biological activities.

Although exogenously applied SLs inhibit shoot branching, the genuine shoot branching inhibitor *in planta* remains elusive.^4,5)^ Recent reports suggest that exuded canonical SLs are not endogenous shoot branching inhibitors; tomato *cyp722c* mutant and rice *Os900* mutant lacking orobanchol and 4-deoxyorobanchol, respectively, do not show increased shoot branching phenotype.^24,25)^ Shunrai exude lesser amounts of SLs and show lower shoot branching number as compared to European cultivars, suggesting that Shunrai produces lesser amounts of SLs acting as rhizospheric signals but higher amounts of SLs functioning as endogenous shoot branching inhibitor(s). Accordingly, Shunrai would be a good material to collect elusive shoot branching inhibitor(s) and to understand the regulation mechanism of shoot branching.

Production and exudation of SLs are strongly regulated by nutrient status.^26)^ In general, phosphate deficiency enhances SL exudation in mycotrophic plants and it may vary with plant species whether nitrogen is also influential in SL exudation. The leguminous plant red clover (*Trifolium pratense* L.) did not respond to nitrogen deficiency as red clover has symbiotic relationship with *Rhizobia* and obtains nitrogen from root nodules.^21)^ By contrast, in the case of non-leguminous plant sorghum [*Sorghum bicolor* (L.) Moench] phosphate deficiency as well as nitrogen deficiency significantly enhanced SL exudation,^19)^ suggesting that sorghum depends on AM symbionts for the supply of both phosphate and nitrogen. However, non-leguminous plant tomato (*Solanum lycopersium* L.) did not enhance SL exudation and the leguminous plant Chinese milk vetch (*Astragalus sinicus* L.) significantly enhanced SL exudation under nitrogen deficiency.^15)^ Therefore, precise regulation mechanisms of SL production and exudation under nutrient deficiency remain elusive.

Both *P. ramosa* and *O. minor* have a relatively wide host range and parasitize various plant species. *P. ramosa* is one of the worst root parasitic weeds infecting major dicot crops including tomato, carrot, legumes, oilseed rape and hemp in both developing and developed countries.^27,28)^ *O. minor* is a federally listed noxious weed in the United States and caused some damages to clover production in Oregon.^10)^ In Japan, *O. minor* was first recorded in 1937 in Chiba prefecture and now is widely distributed throughout Japan except for the cool Hokkaido region.^29)^ So far, more weedy broomrapes, such as *P. ramosa* and *P. aegyptiac*a have not yet invaded Japan but these root parasitic weeds are potential threads to agriculture in Japan.

The weedy broomrapes are difficult to control with herbicides as the parasites obtain water and nutrients from host through the haustorium, a specialized organ which connects to host vasculature.^27)^ Therefore, host-selective herbicides due to rapid detoxification in the host plants are not effective to the parasites attached to the host roots; only less toxic metabolites would enter the parasites through the haustorium. This is one of reasons why broomrapes are causing sever damages to agricultural production in developed countries. Under these circumstances, growing trap crops appears to be a good strategy to mitigate infection of broomrapes and to reduce their seed bank.^8-10)^ Barley as well as wheat is a winter crop and can induces ‘suicidal’ germination of broomrape seeds from autumn to early summer. The present study suggested two European cultivars are good potential trap crops for broomrapes as they strongly induce seed germination of broomrapes and also promote AM colonization. However, further study is needed to develop screening methods for effective trap crops and practical application methods in the fields.

## Acknowledgements

The authors would like to thank emeritus Prof. Koichi Yoneyama (Utsunomiya University) for valuable discussions. This work was supported by Japan Science and Technology Agency (PRESTO, JPMJPR17QA and FOREST, JPMJFR220F) and the Japan Society for the Promotion of Science (KAKENHI 19KK0395).

## Supplementary data

**Fig. S1**. Germination stimulation activities of BL1 and BL2 to broomrapes. *n* = 9-12.

## References

1) C. Parker: Observations on the current status of Orobanche and Striga problems worldwide. Pest Manag. Sci. 65, 453–459 (2009).

2) K. Yoneyama and P. B. Brewer: Strigolactones, how are they synthesized to regulate plant growth and development? Curr. Opin. Plant Biol. 63, 102072 (2021).

3) K. Akiyama, K. Matsuzaki and H. Hayashi: Plant sesquiterpenes induce hyphal branching in arbuscular mycorrhizal fungi. Nature 435, 824–827 (2005).

4) V. Gomez-Roldan, S. Fermas, P. B. Brewer, V. Puech-Pages, E. A. Dun, J. P. Pillot, F. Letisse, R. Matusova, S. Danoun, J. C. Portais, H. Bouwmeester, G. Becard, C. A. Beveridge, C. Rameau and S. F. Rochange: Strigolactone inhibition of shoot branching. Nature 455, 189–194 (2008).

5) M. Umehara, A. Hanada, S. Yoshida, K. Akiyama, T. Arite, N. Takeda-Kamiya, H. Magome, Y. Kamiya, K. Shirasu, K. Yoneyama, J. Kyozuka and S. Yamaguchi: Inhibition of shoot branching by new terpenoid plant hormones. Nature 455, 195–200 (2008).

6) S. Yoshida, H. Kameoka, M. Tempo, K. Akiyama, M. Umehara, S. Yamaguchi, H. Hayashi, J. Kyozuka and K. Shirasu: The D3 F-box protein is a key component in host strigolactone responses essential for arbuscular mycorrhizal symbiosis. New Phytol. 196, 1208–1216 (2012).

7) M. Marzec, A. Daszkowska-Golec, A. Collin, M. Melzer, K. Eggert and I. Szarejko: Barley strigolactone signalling mutant hvd14.d reveals the role of strigolactones in abscisic acid-dependent response to drought. Plant Cell Environ. 43, 2239–2253 (2020).

8) G. Brun, L. Braem, S. Thoiron, K. Gevaert, S. Goormachtig and P. Delavault: Seed germination in parasitic plants: what insights can we expect from strigolactone research? J. Exp. Bot. 69, 2265–2280 (2018).

9) D. Rubiales, M. FernÁNdez-Aparicio, K. Wegmann and D. M. Joel: Revisiting strategies for reducing the seedbank of Orobanche and Phelipanche spp. Weed Res. 49, 23–33 (2009).

10) R. D. Lins, J. B. Colquhoun and C. A. Mallory-Smith: Investigation of wheat as a trap crop for control of Orobanche minor. Weed Res. 46, 313–318 (2006).

11) K. M. Isah and S. T. O. Lagoke: Effects of rotation of trap crop varieties on the reaction of some cereal host crops to Striga hermonthica biotypes. Afr. J. Microbiol. Res. 7, 488–497 (2013).

12) X. Ye, M. Zhang, M. Zhang and Y. Ma: assessing the performance of maize (Zea mays L.) as trap crops for the management of sunflower broomrape (Orobanche cumana Wallr.). Agronomy 10 (2020).

13) T. V. Charnikhova, K. Gaus, A. Lumbroso, M. Sanders, J.-P. Vincken, A. De Mesmaeker, C. P. Ruyter-Spira, C. Screpanti and H. J. Bouwmeester: Zeapyranolactone − A novel strigolactone from maize. Phytochem. Lett. 24, 172–178 (2018).

14) X. Xie, T. Kisugi, K. Yoneyama, T. Nomura, K. Akiyama, K. Uchida, T. Yokota, C. S. P. McErlean and K. Yoneyama: Methyl zealactonoate, a novel germination stimulant for root parasitic weeds produced by maize. J. Pestic. Sci. 42, 58–61 (2017).

15) K. Yoneyama, X. Xie, H. I. Kim, T. Kisugi, T. Nomura, H. Sekimoto, T. Yokota and K. Yoneyama: How do nitrogen and phosphorus deficiencies affect strigolactone production and exudation? Planta 235, 1197–1207 (2012).

16) C. Li, L. Dong, J. Durairaj, J. C. Guan, M. Yoshimura, P. Quinodoz, R. Horber, K. Gaus, J. Li, Y. B. Setotaw, J. Qi, H. De Groote, Y. Wang, B. Thiombiano, K. Flokova, A. Walmsley, T. V. Charnikhova, A. Chojnacka, S. Correia de Lemos, Y. Ding, D. Skibbe, K. Hermann, C. Screpanti, A. De Mesmaeker, E. A. Schmelz, A. Menkir, M. Medema, A. D. J. Van Dijk, J. Wu, K. E. Koch and H. J. Bouwmeester: Maize resistance to witchweed through changes in strigolactone biosynthesis. Science 379, 94–99 (2023).

17) D. Moriyama, T. Wakabayashi, N. Shiotani, S. Yamamoto, Y. Furusato, K. Yabe, M. Mizutani, H. Takikawa and Y. Sugimoto: Identification of 6-epi-heliolactone as a biosynthetic precursor of avenaol in Avena strigosa. Biosci. Biotechnol. Biochem. 86, 998–1003 (2022).

18) N. Arai, K. Yoneyama, M. Inoue, K. Uchida, K. Akiyama and X. Xie, Abstr. 2022 Annu. Meeting Jpn. Soc. Biosci. Biotechnol. Agrochem. 2E05–03 (2022) (in Japanese).

19) K. Yoneyama, X. Xie, D. Kusumoto, H. Sekimoto, Y. Sugimoto, Y. Takeuchi and K. Yoneyama: Nitrogen deficiency as well as phosphorus deficiency in sorghum promotes the production and exudation of 5-deoxystrigol, the host recognition signal for arbuscular mycorrhizal fungi and root parasites. Planta 227, 125–132 (2007).

20) Y. Kobae and R. Ohtomo: An improved method for bright-field imaging of arbuscular mycorrhizal fungi in plant roots. Soil Sci. Plant Nutri. 62, 27–30 (2015).

21) K. Yoneyama, K. Yoneyama, Y. Takeuchi and H. Sekimoto: Phosphorus deficiency in red clover promotes exudation of orobanchol, the signal for mycorrhizal symbionts and germination stimulant for root parasites. Planta 225, 1031–1038 (2007).

22) H. Wang, W. Chen, K. Eggert, T. Charnikhova, H. Bouwmeester, P. Schweizer, M. R. Hajirezaei, C. Seiler, N. Sreenivasulu, N. von Wiren and M. Kuhlmann: Abscisic acid influences tillering by modulation of strigolactones in barley. J. Exp. Bot. 69, 3883–3898 (2018).

23) V. Lailheugue, I. Merlin, S. Boutet, F. Perreau, J. B. Pouvreau, S. Delgrange, P. H. Ducrot, B. Cottyn-Boitte, G. Mouille and V. Lauvergeat: Vitislactone, a non-canonical strigolactone exudated by grapevine rootstocks in response to nitrogen starvation. Phytochemistry 215, 113837 (2023).

24) T. Wakabayashi, M. Hamana, A. Mori, R. Akiyama, K. Ueno, K. Osakabe, Y. Osakabe, H. Suzuki, H. Takikawa, M. Mizutani and Y. Sugimoto: Direct conversion of carlactonoic acid to orobanchol by cytochrome P450 CYP722C in strigolactone biosynthesis. Sci. Adv. 5, eaax9067 (2019).

25) S. Ito, J. Braguy, J. Y. Wang, A. Yoda, V. Fiorilli, I. Takahashi, M. Jamil, A. Felemban, S. Miyazaki, T. Mazzarella, G. E. Chen, A. Shinozawa, A. Balakrishna, L. Berqdar, C. Rajan, S. Ali, I. Haider, Y. Sasaki, S. Yajima, K. Akiyama, L. Lanfranco, M. D. Zurbriggen, T. Nomura, T. Asami and S. Al-Babili: Canonical strigolactones are not the major determinant of tillering but important rhizospheric signals in rice. Sci. Adv. 8, eadd1278 (2022).

26) K. Yoneyama: How Do Strigolactones Ameliorate Nutrient Deficiencies in Plants? Cold Spring Harb Perspect Biol 11 (2019).

27) M. Vurro: Are root parasitic broomrapes still a good target for bioherbicide control? Pest Manag. Sci. 80, 10–18 (2024).

28) A. Casadesus and S. Munne-Bosch: Holoparasitic plant-host interactions and their impact on Mediterranean ecosystems. Plant Physiol. 185, 1325–1338 (2021).

29) Y. Takeuchi: Biology of parasitic weeds and their control. Japanese J. Pestic. Sci. 19, 183–195 (1994) (in Japanese).

