## Supplementary figures and images for "Barley is a potential trap crop for root parasitic weeds broomrapes"

### Supplementary Figure 1

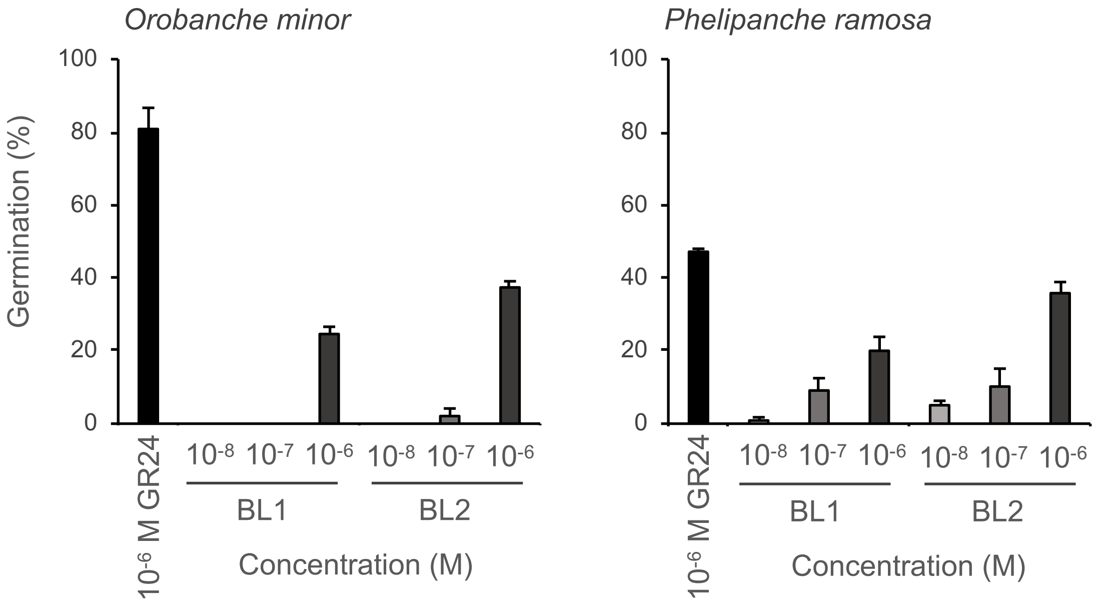
